# Virion aggregation shapes infection dynamics and evolutionary potential

**DOI:** 10.1101/2025.07.03.662980

**Authors:** Meher Sethi, David VanInsberghe, Bernardo A. Mainou, Anice C. Lowen

## Abstract

Viral spread is classically thought to be mediated by single viral particles. However, viruses often disseminate in groups - as aggregates, inside membranous vesicles, or as clusters associated with bacterial components or other complex surfaces. The implications of collective dispersal for viral infectivity and evolution remain incompletely defined. Here, we used mammalian orthoreovirus to evaluate the impact of aggregation on the propagation of infection and the generation of viral diversity through reassortment. Aggregation of free virions was induced by manipulating pH and ionic conditions. This treatment promoted coordinated delivery of viruses to cells, increasing the number of virions per infected cell and number of virions per occupied endosome at early times of infection. Likely due to a consolidation of infectious units, aggregation concomitantly reduced the overall infectivity of the viral population and progeny virus yields. When viral populations comprised two genetically distinct viruses, aggregation increased the frequency of mixed infection and of genetic exchange through reassortment. Thus, the formation of collective infectious units lowers the replicative potential of mammalian orthoreovirus populations but increases viral evolutionary potential by promoting genetic diversification.

**Importance:** A deeper understanding of processes shaping viral evolution will advance our ability to anticipate viral emergence, escape from immune responses and resistance to therapeutics. Although much is known about how genetic variation fuels viral evolution, how modes of viral spread influence the generation and structure of genetic variation remains poorly characterized. Here we examine how the collective dissemination of viruses modulates early infection dynamics and viral diversity. We find that, although infection in groups reduces the number of independently infected cells, it results in a more genetically diverse progeny population, an outcome that may enhance evolutionary potential.

## Introduction

Classical models of viral infection suggest that viruses disseminate and infect cells as individual particles. Many viruses can be transmitted in groups, however, a process known as collective dissemination (1). This process includes both within-host and between-host transmission, enabling simultaneous delivery of multiple viral particles to a single cell or a single host, respectively. Mechanisms of collective dissemination include cell to cell spread via syncytia or cellular junctions such as plasmodesmata, synapses and tunneling nanotubes (2, 3); virion aggregation (4–9); transport of virions within extracellular vesicles (10–14); and scaffolding of virions onto bacterial cells (15–20).

Group-based transmission can confer several advantages over singular infections (21–23). These include enhanced infectivity and more rapid establishment of initial infection (10, 24, 25). For vesicular stomatitis virus (VSV), aggregation increases early-stage reproductive success by raising cellular multiplicity of infection, thereby reducing stochastic infection failure (26). Collective dissemination can also enhance immune evasion by shielding viruses from detection by the host’s immune system (27) and boost viral output through sharing of viral resources (25, 28). In addition, collective infectious units have elevated stability in extracellular environments compared to single particles (29–34). Collective dissemination can also impact the viral evolutionary potential, for example, by enhancing the diversification of viral populations through recombination (16) and by facilitating accumulation of defective interfering (DI) particles over multiple generations (37).

Mammalian orthoreovirus (reovirus), a member of the *Reoviridae*, is a non-enveloped virus with a segmented double stranded (ds) RNA genome comprising 10 segments (38, 39). Reovirus primarily infects epithelial cells of the gastrointestinal and respiratory tracts, but can also infect the heart and central nervous system in human infants and murine models (40–45). Reovirus enters cells via receptor-mediated endocytosis, followed by partial disassembly in endosomes to release transcriptionally active cores into the cytoplasm where replication initiates inside non-membranous structures called inclusion bodies (46, 47).

For viruses with segmented genomes, co-infection enables reassortment, a form of genetic exchange in which intact genome segments are shuffled into novel combinations (48–51). Reassortment can be a major source of viral diversity: a cell co-infected with two reoviruses that differ in all ten segments can generate 2^10^ (1024) unique reassortants. In line with this theoretical prediction, we and others found that reovirus reassortment occurs with high efficiency in co-infected cells (52, 53). This high reassortment capacity makes reovirus an ideal model for investigating how collective dissemination and co-infection influence reassortment and viral diversity.

Virus aggregation offers a tractable and physiologically relevant system for studying collective dissemination. Reovirus engages in collective spread via adhesion to bacteria or dispersal inside extracellular vesicles (12, 29, 54). Reovirus also aggregates near its isoelectric point, where surface charge is minimized and interparticle attraction is favored (55–57). As an enteric virus, reovirus encounters dramatic pH gradients (∼1,5 to ∼7.4) during transit through the gastrointestinal tract (58). While the impact of these pH fluctuations on aggregation *in vivo* remains unclear, acidic conditions have been shown to induce aggregation *in vitro* (59, 60). This aggregation is reversible and can be modulated by adjusting environmental pH, enabling experimental control (7).

Here, we used reovirus aggregation to test our hypothesis that collective dispersal promotes genetic diversification through reassortment. In parallel, we examined whether aggregation limits viral spread by consolidating infectious units. We manipulated viral aggregation state using low pH and assessed outcomes through co-infection assays, genotyping viral progeny, and infectivity measurements. We find that collective dissemination augments viral diversity but that this potential evolutionary benefit comes at a cost of reduced replication efficiency.

## Results

### Reovirus aggregation is pH dependent, reversible and influenced by buffer composition

To probe the evolutionary implications of collective dissemination, we established a system for modulating the degree of aggregation of mammalian orthoreovirus type 3 Dearing (T3D). We leveraged the work of Floyd and Sharp, who showed that virion aggregation is triggered by changes in pH and ionic conditions and that reovirus aggregates around its isoelectric point of pH 3.9 (7, 55, 59, 61). Guided by their findings, we performed a targeted screen of buffers ranging from pH 4 to pH 7.4, analyzing virion aggregation by transmission electron microscopy (TEM) **(Fig 1)**. We found that, when the virus was suspended in PBS or incubated in acetate pH 5 buffer, most viral objects comprised five or fewer virions with a predominance of single, unaggregated virions. Incubation in tris citrate pH 5 gave an intermediate outcome and incubation in any of the three pH 4 buffers (tris-citrate, acetate and citrate) yielded a substantial increase in the frequency of larger aggregates **(Fig 1A, 1B)**. Given the prominent aggregation observed in acetate and tris-citrate pH 4 buffers, additional replicates were analyzed with these conditions to confirm reproducibility and identify aggregation conditions suitable for downstream studies **(Fig 1C)**.

**Figure 1:**
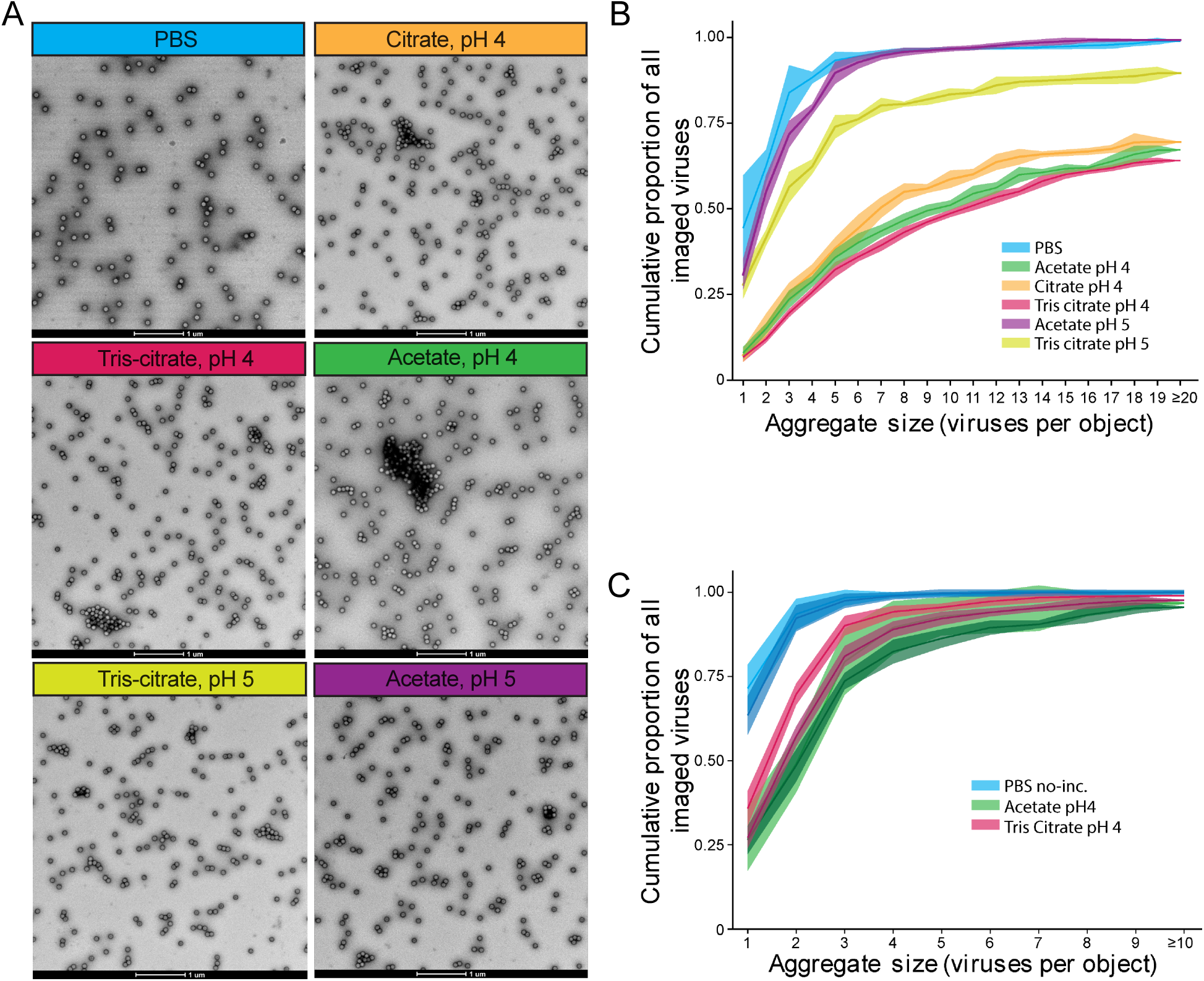
Reovirus aggregation is enhanced near the isoelectric point and further tuned by ionic properties of the buffer. **(A)** Representative transmission electron microscopy (TEM) images of purified T3D reovirus across 6 different buffer conditions. Scale bars, 1 μm. **(B)** Cumulative proportion of the number of virions per object across six different buffer conditions. N=1 per condition. The mean and 95% CI derived from 16 fields per condition are plotted. **(C)** Quantification of aggregation levels in pH 4 buffers was measured by DLS, based on the size of viral objects. Data were pooled from 3 biological replicates. Each data point represents a viral object of size ≥ 1. Bars represent the geometric mean. *p < 0.05; ns, non-significant by Kruskal-Walli test followed by Dunn’s multiple comparison test, with false discovery rate controlled using the two-stage step-up method of Benjamini, Krieger, and Yekutieli.

For downstream experiments, it was important to consider the impact of buffer composition on cell health (62). L929 cells showed a significant reduction in viability following exposure to PBS and all low pH buffers tested (**Supp fig 1A**). To mitigate cytotoxicity, we diluted these buffers in Opti-MEM. We initially diluted the non-aggregated inoculum (PBS) by 50% and aggregated inoculums (acetate, tris-citrate and citrate) to a composition of 30% low pH buffer and 70% Opti-MEM. However, dilution of the low pH buffers in Opti-MEM reduced the extent of viral aggregation (**Supp fig 1B**). We then tested a broader range of conditions. Except for acetate, dilution to 70% buffer and 30% Opti-MEM improved cell viability (**Supp fig 2A**). Selecting tris-citrate pH 4 for further examination, we evaluated the impact of dilution on aggregation by dynamic light scattering (DLS) (a complimentary approach to TEM image analysis that is suitable for more dilute samples) **(Supp fig 2B)**. Aggregation levels remained elevated across all tris-citrate formulations compared to PBS, and the 70% formulation resulting in larger aggregates than the 50% condition **(Supp fig 2B)**.

Finally, we sought to confirm that the tris-citrate-Opti-MEM mixtures do not impair viral function. We first assessed infectivity in buffer in the absence of aggregation, by omitting the incubation of virus in the buffer. Viral replication and infectivity were comparable between the tris citrate and PBS control inoculums **(Supp fig 2C, 2D).** We then assessed whether reovirus infectivity is diminished by incubation at low pH. Reovirus was first aggregated using low pH buffer (tris-citrate, pH 4) and aggregation was then reversed by dilution to a final formulation of 15% tris-citrate in Opti-MEM. Reversal of aggregation was confirmed using DLS **(Supp Figure 3A)** and viral infectivity was assessed by plaque assay **(Supp fig 3B)**. Aggregated samples showed reduced titers, as expected, but reversal of aggregation restored infectivity to levels comparable to non-aggregated virus prepared in PBS. Thus, infectivity of T3D reovirus particles was not compromised by incubation in tris-citrate, pH 4 buffer.

For downstream experiments investigating collective dissemination, non-aggregated virus was prepared by suspending virus in a 1:1 mixture of PBS and Opti-MEM and aggregated virus was prepared through incubation in tris-citrate pH 4, followed by addition of Opti-MEM prior to infection, such that tris-citrate made up 50% or 70% of the final inoculum volume.

### Aggregation ensures coordinated delivery of viruses to cells

To evaluate whether viral aggregation enhanced multi-virion delivery to individual cells, we infected cells with virus prepared under non-aggregating and aggregating conditions. We used TEM to determine if clusters remain intact during viral entry **(Fig 2)**, if aggregation increases the likelihood of multiple viruses being taken up into endosomes **(Fig 3)**, and if aggregation results in greater intracellular virion accumulation per cell **(Fig 4)**.

**Figure 2:**
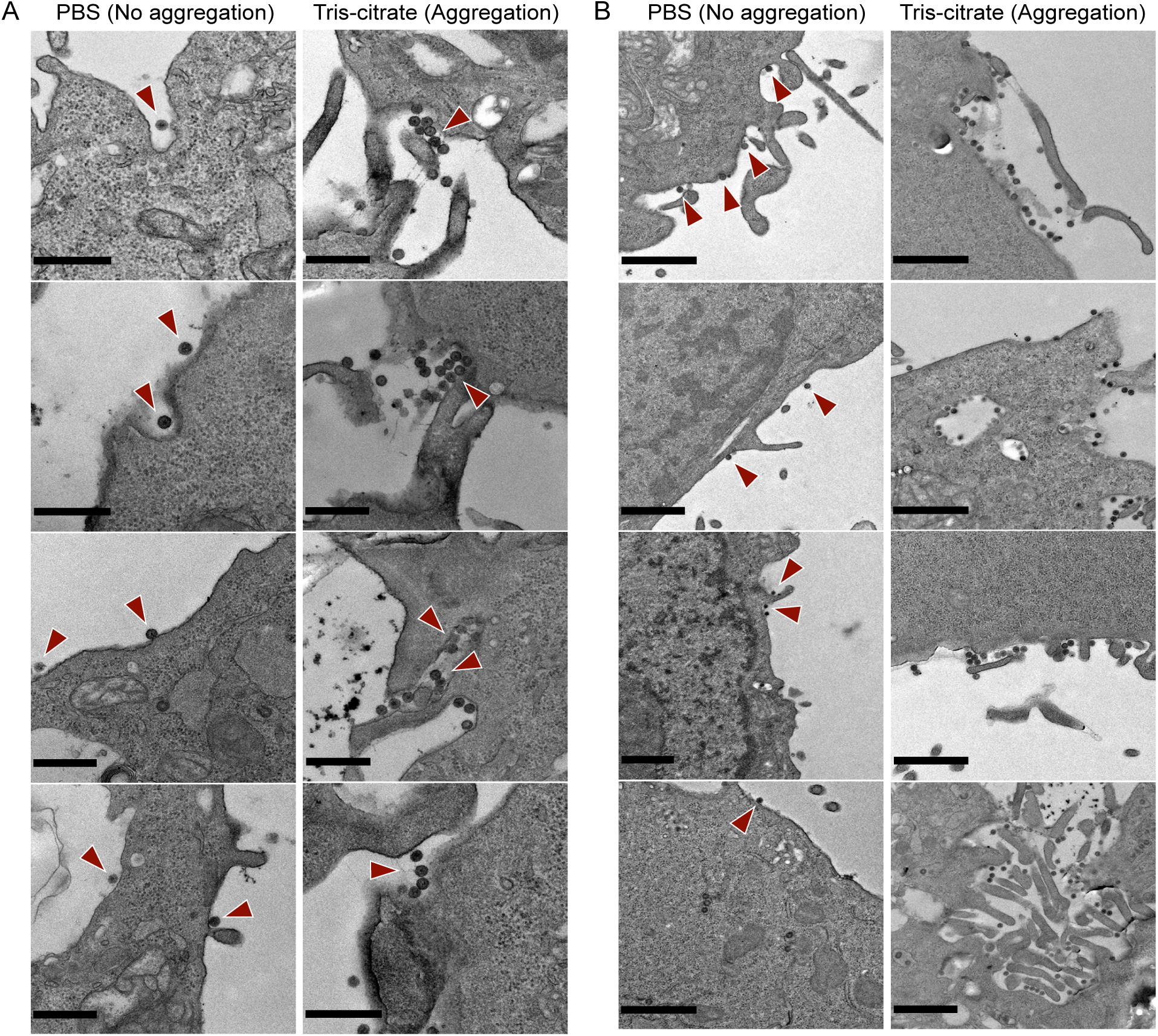
Collective entry of reovirus aggregates into cells. **(A-B)** Representative TEM images of L929 cells infected with either non-aggregated or aggregated reovirus particles. Cells were infected at an MOI of 10 PFU/mL and allowed to adsorb for 5 minutes before sample processing. **(A)** Aggregated virions are shown entering cells as clusters. Scale bars, 1 μm. Red arrowheads highlight single virions (PBS) and aggregated viruses (Tris-citrate) entering the cell. **(B)** Dispersed viral aggregates give rise to concentrated patches of viruses at the cell surface. Red arrowheads (in PBS) highlight individual virions at the cell surface and illustrate typical morphology and appearance of virions. Scale bars represent 1 μm.

**Figure 3:**
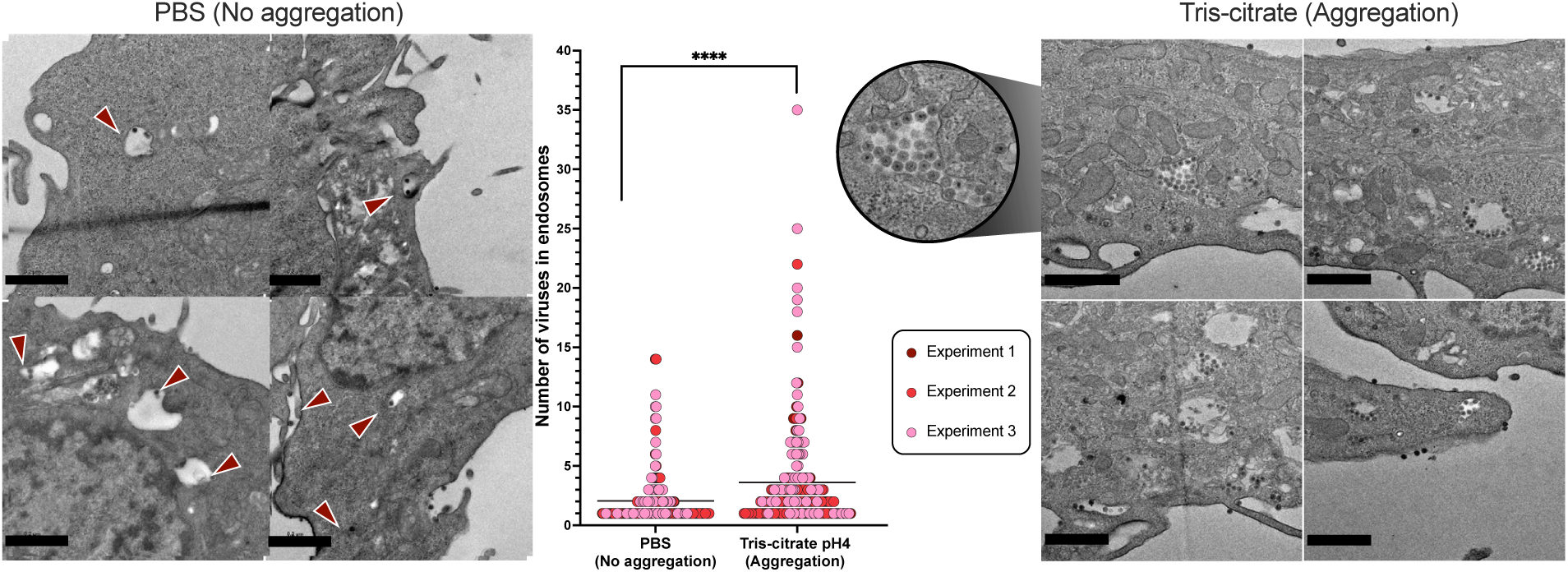
Aggregation promotes multi-virion delivery to individual cells. Representative TEM images of L929 cells infected with either non-aggregated (left) or aggregated (right) reovirus particles. Cells were inoculated with 10 PFU/cell and virus was allowed to adsorb for 60 minutes before sample processing. Endosomes containing multiple viruses are visible in cells infected with aggregated viruses, whereas cells infected with non-aggregated virus primarily contain ≤ 2 virions within the endosome. Inset from the aggregation condition shows a magnified endosome densely packed with virions. Plot at center shows the number of viruses per endosome in cells infected with non-aggregated and aggregated virions. Data are pooled from three experiments. Each experiment is represented by a distinct color, and each data point corresponds to the number of virions counted within a single endosome. Horizontal lines indicate means. ****p<0.0001, unpaired Mann-Whitney test. Scale bars, 1 μm.

**Figure 4:**
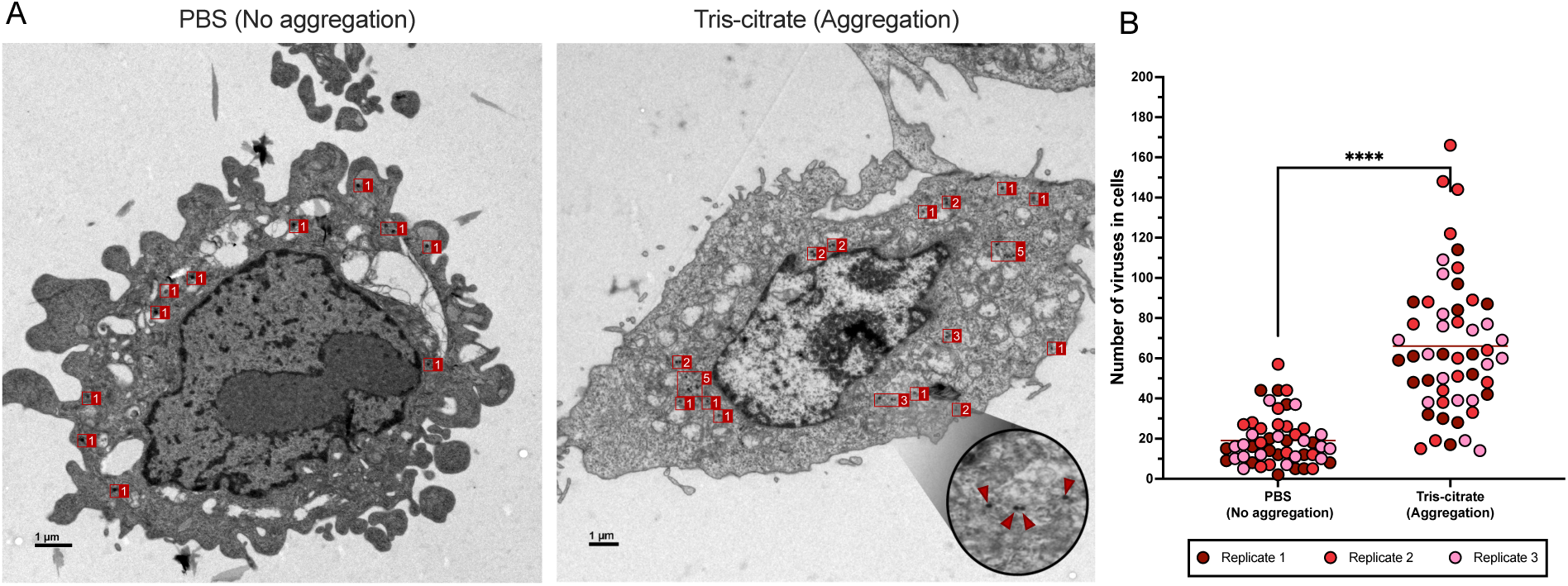
Aggregation increases the number of virions per infected cell. **(A)** Representative TEM images of L929 cells infected with either non-aggregated or aggregated reovirus particles. Cells were inoculated with 10 PFU/cell and virus was allowed to adsorb for 10 minutes prior to sample processing. Red boxes mark virions; adjacent numbers indicate the count of virions within each box. Inset from the aggregation condition shows a magnified region of the cell highlighting the appearance of individual virions (red arrowheads). Scale bars, 1 μm. **(B)** Quantification of intracellular viral particles for aggregated and non-aggregated conditions. Data is pooled from 3 experiments. Each experiment is represented by a distinct color, and each data point corresponds to the number of virions counted within a cell. The horizontal line indicates the grand mean. p<0.0001 by unpaired Mann-Whitney test.

In cells infected with viral aggregates, large clusters of virions were frequently observed at the cell surface entering as a group **(Fig 2A)**. In some instances, aggregates appeared to dissociate at the cell membrane, such that virions entered in patches along the cell surface **(Fig 2B)**. In contrast, cells infected with non-aggregated virus showed individual virions sparsely distributed along the membrane **(Fig 2)**. Within infected cells, individual endosomes were observed to carry multiple virions more frequently when virus was aggregated compared to when it was not **(Fig 3)**. Finally, infected cells treated with aggregated virus showed a markedly higher number of intracellular virions per cell compared to those infected with non-aggregated virus **(Fig 4)**. Together, our TEM analysis demonstrates that viral aggregation facilitates simultaneous entry and uptake of multiple viral particles within individual cells.

### Aggregation promotes co-infection and genetic exchange

To evaluate whether aggregation enhances reassortment, we assessed the frequencies of cellular co-infection and genetic exchange between wild-type (WT) and Variant (Var) parental viruses. Var differs from WT by a single synonymous mutation per gene segment, enabling its differentiation from WT but ensuring comparable fitness. The use of WT and Var therefore allows quantification of co-infection and reassortment without bias resulting from differing replication kinetics of the two viruses (52, 63). We prepared a 1:1 mixture of WT and Var viruses and validated the ratio using two approaches: quantification of WT and Var genome copies (GC) within the bulk mixture and genotyping of 48 plaque isolates derived from the mixture. Both methods showed comparable representation of WT and Var, confirming a 1:1 mixture **(Supp fig 4A-B)**.

Using the 1:1 WT-Var inoculum, we measured the impact of aggregation on the frequency of cellular co-infection. To this end, we treated the 1:1 mixture under aggregating and non-aggregating conditions and then isolated 32 plaques from each preparation from three independent experiments. Plaques were genotyped to determine whether the viruses therein were WT-only, Var-only, or mixed. We found that aggregated samples showed a significantly higher proportion of mixed plaques, indicating that aggregation promotes co-infection **(Fig 5A)**.

**Figure 5:**
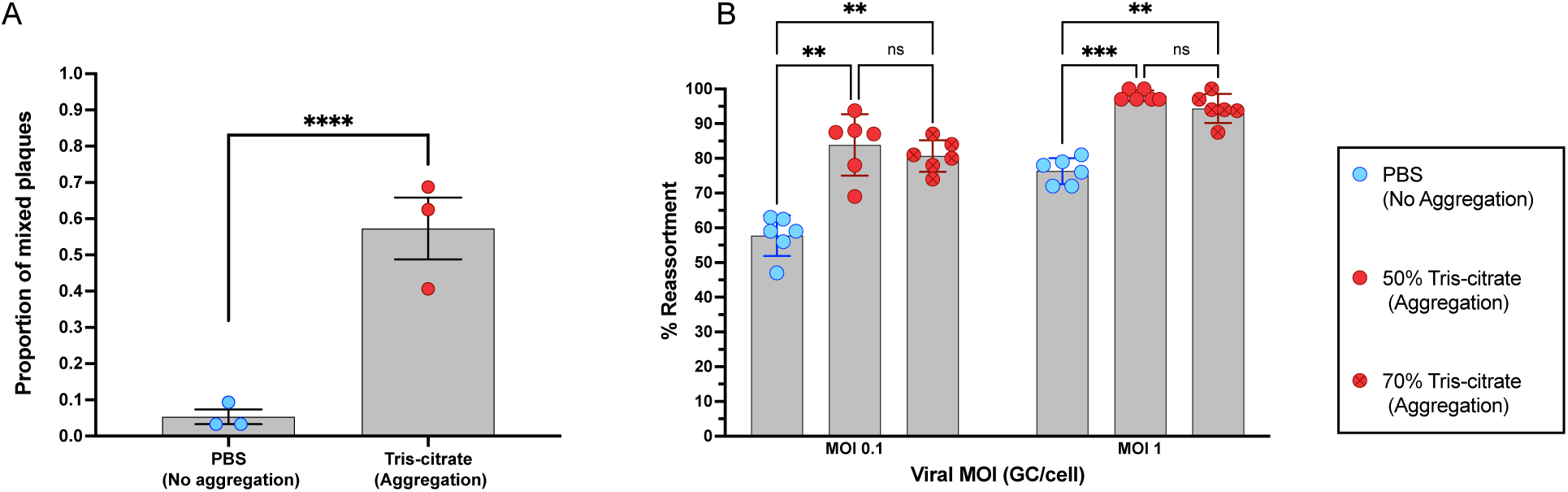
Aggregation promotes co-infection and enhances reassortment. **(A)** Genotypes of viral plaques were determined following infection of L929 cells with aggregated (50% tris-citrate in Opti-MEM) or non-aggregated (50% PBS in Opti-MEM) inoculum of WT and Var reoviruses at a multiplicity of 0.1 GC/cell. Mixed plaques were positive for both WT and Var genomes. Data shown are mean ± s.d of three biological replicates. Each data point reflects the proportion of mixed plaques among 32 plaques. p<0.0001, unpaired Mann-Whitney test. **(B)** Reassortment frequency was assessed by defining the percentage of plaque isolates with any reassortant genotype, following multicycle replication in cells co-infected with WT and Var reoviruses. Data shown are mean ± s.d. and represent 6 biological replicates derived from two independent experiments. Each data point represents the percentage of reassortant plaques out of 32 plaques. ***p<0.001; **p<0.01; ns, not significant, two-way ANOVA followed by Sidak’s multiple comparison test.

We next examined how aggregation modulates the frequency of reassortment. Cells were co-infected with WT and Var using the 1:1 viral inoculum at MOIs of 0.1 or 1 GC/cell, and multicycle replication was allowed to proceed. Levels of reassortment were determined by analyzing progeny from six biological replicates, genotyping 32 plaque isolates per replicate. We observed that both aggregation formulations (50% and 70% tris-citrate) showed increased frequencies of reassortant progeny compared to the non-aggregated control at both MOIs **(Fig 5B)**. To confirm that the observed effects were not an artifact of buffer composition, we validated PBS as a reliable negative control by comparing virus prepared in PBS to that made in tris-citrate buffer without incubation **(Supp fig 4C)**. As expected, when the incubation period that allows virion aggregation was omitted, no significant differences in reassortment frequencies were observed among inoculums delivered in PBS and 50% and 70% tris-citrate formulations **(Supp fig 4C)**. Together, these findings indicate that viral aggregation increases the likelihood of co-infection, which in turn facilitates genetic exchange through reassortment.

### Aggregation modulates infection dynamics

To test the impact of aggregation on viral infection dynamics, we first evaluated the infectious potential of the inoculum. To this end, we performed plaque assays using virus preparations made under aggregating and non-aggregating conditions. Since virion concentration could affect aggregation efficiency, preparations the mimicked those used for infections at MOI 0.1 and 1 GC/cell were tested. Both aggregation conditions (50% and 70% tris-citrate) yielded significantly lower titers compared to non-aggregated virus (PBS) **(Fig 6A)**. Additionally, the 70% tris-citrate condition, which showed a higher frequency of large aggregates **(Supp fig 2B)**, was associated with slightly lower titers compared to the 50% tris-citrate condition. Next, we assessed viral output following multicycle replication in cultures infected at the two MOIs. Consistent with fewer cells being infected at the time of inoculation, progeny viral titers were significantly reduced in 70% tris-citrate aggregation condition at both MOIs **(Fig 6B-C)**. In the 50% tris-citrate condition, however, progeny viral titers remained comparable with PBS at both MOIs **(Fig 6C),** potentially owing to the opportunity for amplification through multiple rounds of infection. To conclude, while aggregation enhances coordinated delivery of multiple viruses to cells and reassortment, it imposes cost by limiting viral output.

**Figure 6:**
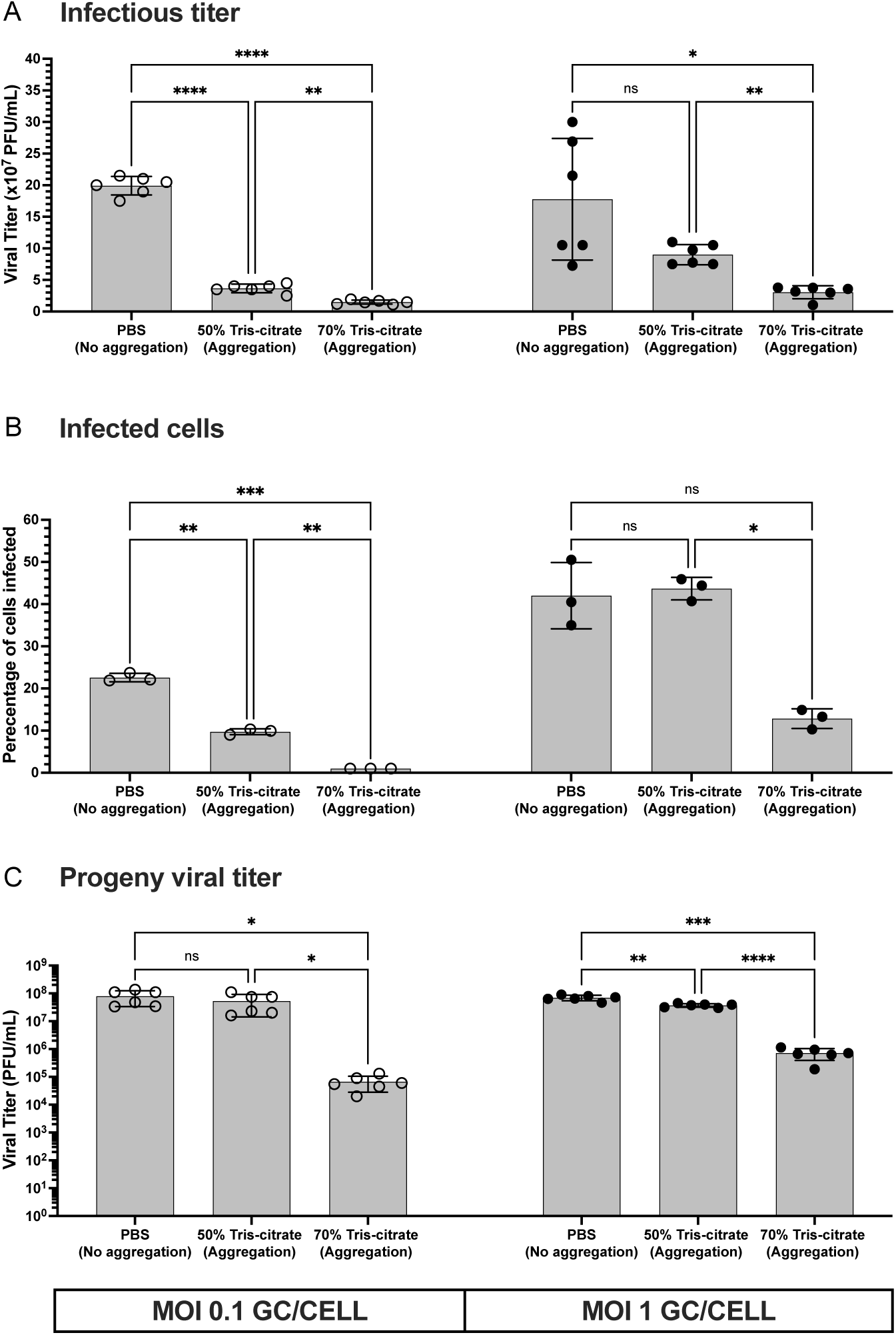
Aggregation limits viral spread and reduces overall viral output. L929 cells were co-infected with non-aggregated virus or virus prepared under one of two aggregation conditions (50% and 70% tris-citrate) at 0.1 or 1 genome copy (GC)/cell. **(A)** Viral titers of the input inoculums. **(B)** The percentage of cells infected at 24 h post inoculation, as quantified by flow cytometry. **(C)** Progeny viral output measured at 24 h post infection. **(A-C)** Each data point represents a biological replicate. Bars represent the mean ± s.d. ****p < 0.0001; ***p < 0.001; **p<0.01; *p<0.05; ns, not significant, two-way ANOVA followed by Tukey’s multiple comparison test.

## Discussion

We investigated how reovirus aggregation, a form of collective dispersal, shapes viral genetic exchange and infection dynamics. We found that aggregation increases the frequency with which multiple viruses enter the same cell and, under conditions of mixed infection, enhances reassortment. However, high levels of aggregation reduced replication efficiency. These findings suggest that collective dispersal can augment viral evolutionary potential but incurs the cost of reduced dissemination.

Several distinct mechanisms of viral clustering can mediate collective dissemination (7, 9, 12, 17). Here, we focus on aggregation governed by direct physiochemical interactions (7, 64). Aggregation likely occurs as the surrounding pH approaches the virus’s isoelectric point, reducing net surface charge and enabling transient particle-particle associations (56, 65). This type of aggregation is dynamic and reversible (7), suggesting non-specific electrostatic interactions may modulate infection efficiency in response to environmental pH. Consistent with prior work (55, 59, 61), we show that exposure of reovirus to low pH near its isoelectric point (∼3.9) promoted aggregation and this aggregation was reversible. Importantly, this mode of aggregation is likely physiologically relevant, as reovirus encounters pH ranging from 1.5 to 7.4 during gastrointestinal transit (55, 58).

Our data show that virion aggregation facilitates group-based delivery, increasing the number of virions delivered to the same cell and resulting in high intracellular virus accumulation. When multiple viral genotypes are present, we found that aggregation enhances the frequency of their co-infection and reassortment, leading to the production of genetically diverse progeny. Diversity produced through reassortment may accelerate the emergence of high fitness variants under selective pressure (66–72). Thus, aggregation may offer a benefit by increasing the adaptability of viral populations. Of note, not all modes of viral clustering are likely to enhance diversity (36). Viral aggregates that form extracellularly can include progeny viruses from distinct foci of infection, which are more likely to differ genetically. In contrast, vesicular ensembles of viruses formed within cells, would be more likely to carry closely related viruses replicated from the same parental genome (36). In the latter case, co-infection and reassortment would not increase diversity (36).

Infection of single cells with multiple viruses can be beneficial for viral replication. For example, the speed and productivity of influenza A virus replication is often increased under conditions conducive to multiple infection, owing to complementation of incomplete viral genomes and increased efficiency of replication under adverse conditions (73–75). While the frequency of incomplete genomes for reovirus is unknown, the production of defective interfering particles that lack portions of essential ORFs has been documented (76). Collective dissemination would allow complementation of defective interfering genomes, but these genomes may then suppress virus production from the cells that carry them (35, 77). We found that, despite increased intracellular viral density, high levels of reovirus aggregation resulted in reduced progeny output. This reduction was likely due to consolidation of infectious units leading to fewer infected cells, an effect that was not fully counterbalanced by any potential benefits of multiple infection. Intermediate levels of aggregation did not result in detectable replication deficits, however, suggesting that costs and benefits of multiple infection may be balanced in this context.

Some limitations of our study are important to consider. First, to allow controlled manipulation of aggregation conditions, the study was conducted in an *in vitro* system, which does not capture the complexity of *in vivo* environments. Aspects of the host environment, such as spatiotemporal variation in pH and the presence of bacteria (15, 29, 17, 7, 65) can modulate viral clustering and may alter the balance of costs and benefits associated with collective dissemination. Second, experimental manipulation of viral aggregation was applied only to the inoculum and did not persist through multiple rounds of viral replication. This factor limits our ability to interpret results quantitatively. Third, we focused on a single reovirus strain, T3D, leaving uncertainty in the extent to which our findings apply across other reovirus strains and viral families.

In conclusion, our results support a model in which virion aggregation mediates collective dissemination, enhancing group-based delivery, co-infection and reassortment under certain environmental conditions, in turn increasing viral population diversity and evolutionary potential. Aggregation shifts viral dynamics from widespread, low multiplicity infection to localized, high multiplicity infection, increasing genetic exchange but limiting dispersal (**Fig 7**). In this way, aggregation may balance short term replication inefficiency with long-term evolutionary resilience.

**Figure 7:**
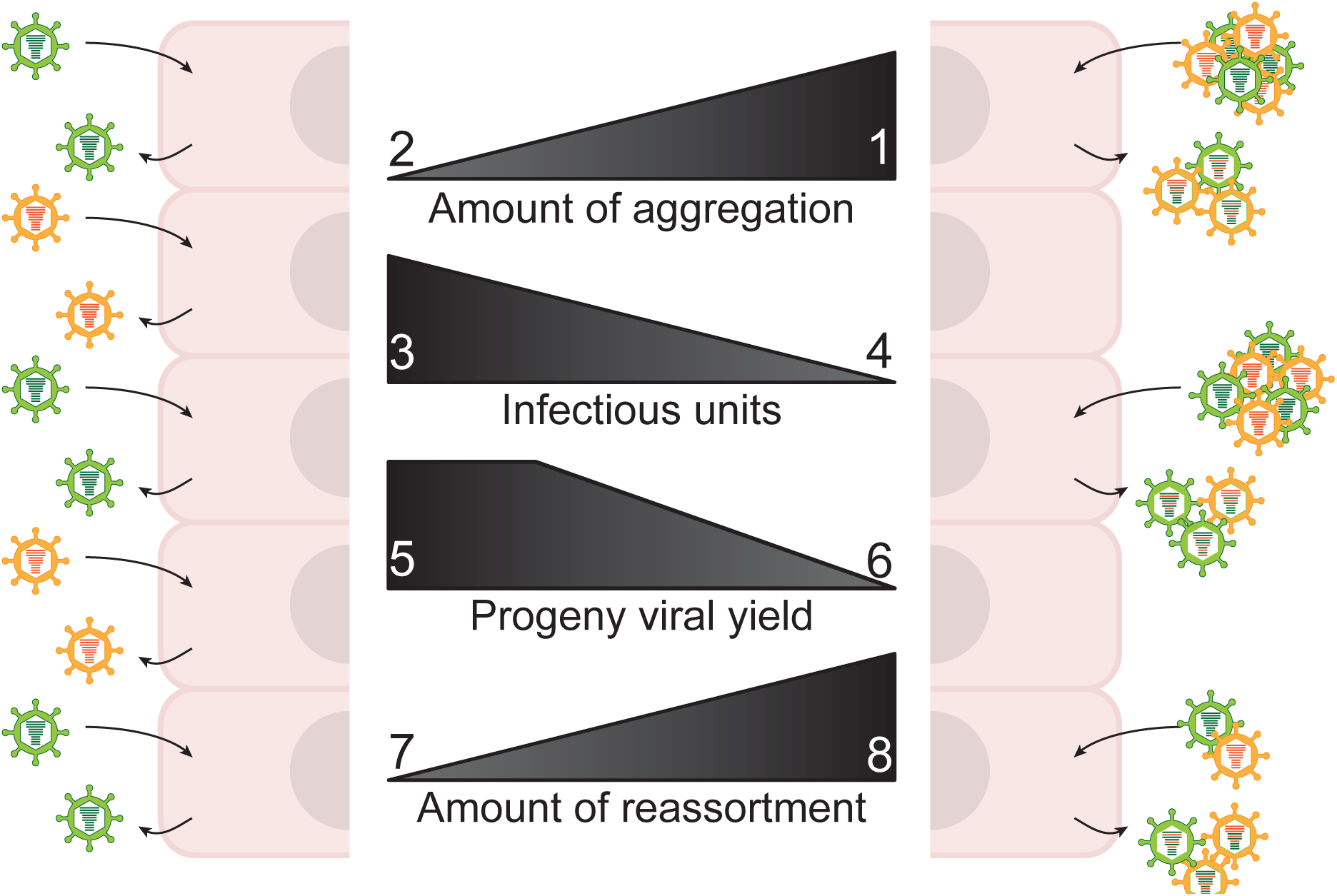
Intercellular virus-virus interactions shape infection dynamics and influence viral evolution, with aggregation limiting spread while enhancing genetic diversity. As the extent of viral aggregation increases (1), multiple virions are delivered to the same cell, reducing the number of independently infected cells and thereby lowering the overall number of infection events (4). At low levels of aggregation (2), viral yield is largely maintained (5). However, under high aggregation conditions where large aggregates are formed (1), yield decreases (6). Importantly, reassortment increases (8) with aggregation due to enhanced co-infection, resulting in greater genetic diversity among progeny and increasing evolutionary potential.

## Materials and Methods

### Cells

Spinner adapted L929 (NCTC clone 929 mouse fibroblast) cells (a gift from Dr. Terry Dermody, University of Pittsburgh) were maintained at 37°C in a humidified atmosphere without CO_2_ supplementation in complete Joklik’s modified eagle medium (JMEM). For experiments, L929 cells were seeded and incubated at 37°C in a humidified atmosphere containing 5% CO_2_, which induced an adherent phenotype. Complete JMEM was prepared from a powdered formulation (US Biological # M3867). The pH was adjusted to 7.2 using 10 N NaOH. The medium was then supplemented with 5% FBS, 2 mM L-glutamine (Corning), 1% penicillin-streptomycin (PS) (Corning), and 0.25 mg/mL amphotericin B (Sigma-Aldrich). Prior to use, the medium was sterilized by filtration through a 0.22 µm polyether sulfone membrane (PES) filter (Corning). BHKT-7 cells (78) were cultured in Dulbecco’s Modified Eagle’s Medium (DMEM) (Gibco) supplemented with 5% FBS, 2 mM L-glutamine, PS, and 1 mg/mL G418 (Invivogen) at 37°C in 5% CO_2_. Cells were tested for mycoplasma contamination monthly, and any positive cultures were promptly discarded.

### Viruses

Wild-type (WT) T3D reovirus (NCBI accession no. SRX6802327, and Variant (Var) (PMID 31511390) T3D reovirus (63) were generated by reverse genetics (79, 80). T3D Var virus contained the following previously described point mutations: L1 C612T, L2 C853T, L3 G481A, M1 C919T, M2 A650G, M3 T702C, S1 G312A, S2 A438G, S3 T318C, and S4 C383T. Both viruses were produced in BHK-T7 cells and propagated in L929 cells for three serial passages. Viral stocks were subjected to Vertrel-XF extraction and the virus in the resulting aqueous fraction was purified by ultracentrifugation through a 1.2-1.4 g/mL cesium chloride gradient. The resulting viral band was collected, dialyzed and stored at 4°C (81).

### Preparation of aggregated and non-aggregated forms of T3D reovirus

To induce aggregation, T3D reovirus was added to low pH buffers at a final concentration of 2.7 x 10^4^ genome copies (GC)/µL. The mixture was then left undisturbed at room temperature (RT) for 4 h to allow aggregation. For the non-aggregated control, virus was re-suspended in 1X PBS and used immediately.

Buffers for aggregation were prepared in cell culture grade water (Corning), filtered through 0.22 µm PES filter (Corning), and stored at 4°C. Tris-citrate buffers (0.05 M) were prepared using 0.1 M Tris base and 0.04 M citric acid. Acetate buffers (0.05 M) were prepared using glacial acetic acid. Citrate buffer (0.05 M) was prepared using citric acid. In each case, the final pH was adjusted using 10 N NaOH.

To minimize L929 cytotoxicity associated with exposure to low pH buffers and PBS (Corning), Opti-MEM (Gibco) was added to viral samples prior to infection. Non-aggregated control virus was resuspended in a 1:1 mixture of PBS and Opti-MEM without incubation. Aggregated preparations were combined with Opti-MEM to yield mixtures in which tris-citrate buffer constituted either 50% or 70% of the total volume. All virus preparations were used for infection immediately following dilution with Opti-MEM.

### Transmission electron microscopy (TEM) to visualize viral aggregates

To analyze virus aggregation, TEM was performed using a Talos 120C microscope. Virus samples of 2.7 x 10^5^ GC in 10 µL of specified buffer (final concentration 2.7 x 10^4^ GC/µL) were prepared for negative staining as follows. A 0.75% (w/v) uranyl formate solution was freshly prepared each day by dissolving 0.0375 g uranyl formate stain in 5 mL of warm de-ionized (DI) water. The solution was vortexed thoroughly to disperse any clumps, followed by the addition of 4 µL of 10 N NaOH and filtered through a 0.22 µm syringe filter. 400mesh copper square grids with 10 nm carbon film (VWR # 103303-424) were glow discharged to increase hydrophilicity and ensure sample adhesion. For each sample, 3 µL of virus suspension was applied to the grid and allowed to absorb for 1 minute before rinsing twice with a drop of ultra-pure water and blotted dry. The grid was then stained with 0.75% (w/v) uranyl formate solution for 1 minute and blot dried before visualization.

### Automated analysis of TEM images

TEM images were processed using a custom python script to detect viral aggregates, followed by a supervised machine learning classifier to determine aggregate sizes. The script uses the python packages scikit-image (82) for image processing and scikit-learn (83) for the classifier. Briefly, images were first processed to remove background noise and intensity variation within images using a rolling-ball filter, then a threshold was set and labeled to identify discrete objects (aggregates of different sizes). Using a script, objects were then randomly selected for manual classification of aggregate sizes, where ideally at least 10 objects per image were manually classified. The classifier was then trained using the area, perimeter, solidity, mean intensity, background intensity, maximum axis, and minimum axis values from the manually classified objects and used to define aggregates.

### Electron microscopy to visualize viral entry

L929 cell monolayers (5×10^5^ cells/well in 24-well plate) were infected with aggregated (50% tris-citrate inoculum) or non-aggregated (1:1 PBS-Opti-MEM mixture) forms of T3D reovirus. Following infection for the designated time, cells were fixed in the culture plate without removing the inoculum, with a mixture of 2.5% glutaraldehyde, 1.0 % paraformaldehyde, 2.6 mM MgCl_2_, 2.6 mM CaCl_2_, 50 mM KCl, 0.01% Picric Acid, and 2% sucrose in 0.1 M cacodylate buffer overnight at 4°C. The fixative was then removed, and cells were washed 3 times with fresh 0.1 M cacodylate buffer at pH 7.4. The cells underwent a second round of fixation with 1.0% osmium tetroxide for 1 h to strengthen the cellular cytoskeleton. The cell monolayers still attached to the culture plate were then washed with water and stained *en bloc* with 2% aqueous uranyl acetate for 20 min at 60°C. Monolayers were then dehydrated in a stepwise manner in 25%, 50%, 70%, and 95% ethanol, followed by three changes in 100% ethanol. Monolayers were then infiltrated with Eponate 12 resin and polymerized for 48 h at 60°C. Polymerized blocks were then removed from the culture plate with the cells flat embedded in the bottom of the block. Ultrathin sections were cut from the bottom of the block with a Leica EM UC6 ultramicrotome, stained with uranyl acetate and lead citrate, and imaged in a Jeol JEM 1400 TEM operated at 80KV. Electron micrographs were acquired on a 2048×2048 charge-coupled device (CCD) camera (UltraScan 1000, Gatan Inc, Pleasanton, CA, USA).

### Cell viability assay to measure cytotoxicity of buffers

L929 cells (1.5×10^4^ cells/well) were seeded in 96-well, clear bottom plates and incubated overnight. After removing culture medium, the cell monolayers were treated with 20 µL of buffer mixed with escalating proportions of Opti-MEM for 1 h on a shaker in a 37°C and 5% CO_2_ incubator. After 1 h, the buffer-Opti-MEM mixtures were removed and 200 µL of complete JMEM was added to each well. Cell viability was tested using the Cell Proliferation Kit (Roche # 11465007001), following manufacturer’s instructions. Briefly, the 96-well culture plates were incubated for an additional 6 h at 37°C and 5% CO_2_. Next, 10 µL of MTT (3-(4,5-Dimethyl-2-thiazolyl)-2,5-diphenyl-2H-tetrazolium Bromide) reagent at a final concentration of 0.5 mg/mL was added to each well and mixed thoroughly. After 5 h, 100 µL of solubilization buffer (10% SDS in 0.01 M HCl) was added to each well to dissolve the purple formazan crystals formed by viable cells. The solubilization reaction was allowed to proceed for 14 h at 37°C and 5% CO_2_ before absorbance was read at a wavelength of 580 nm using a Synergy H1 plate reader (BioTek).

### Dynamic light scattering (DLS) to analyze virus aggregation

To assess particle size and aggregation state, DLS was performed using Wyatt Dynapro instrument with Dynamics 8.3.1 software. Each sample contained 3.5 x 10^8^ GC/mL of virus, resuspended in PBS or incubated in the designated low pH buffer to induce aggregation; Opti-MEM was included where indicated.

For standard DLS experiments measuring aggregation levels, a 200 µL sample was prepared and run in three 10 µL technical replicates using disposable micro cuvettes. For reversal of aggregation, the full 200 µL sample was imaged by splitting the inoculum across three cuvettes.

DLS detects time-dependent fluctuation in scattered light intensity caused by Brownian motion. These fluctuations were used to calculate the diffusion coefficient, and the hydrodynamic radius (Rh) was derived using the Stokes-Einstein equation by the Dynamics 8.3.1 software. Five measurements, each consisting of 10 acquisitions per sample, were used to generate average size distributions.

To estimate virion counts per object, we assumed an average virion diameter of 80 nm and used the following formula.

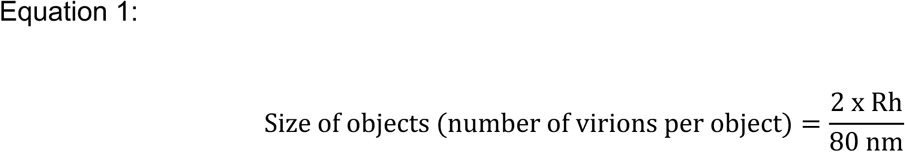

### Droplet digital PCR (ddPCR) to measure genome copies

To prepare samples for ddPCR, total RNA was extracted from 10 µL of viral stocks (WT, Var and 1:1 mixture) using the QIAamp Viral RNA mini kit (Qiagen), following the manufacturer’s protocol (carrier RNA was not added). RNA was eluted in 40 µL of ultra-pure water and stored at -80°C until use. cDNA was synthesized from 12.8 µL of RNA in 20 µL reactions using Maxima reverse transcriptase (RT) (Thermo Scientific, Cat # EP0742) with the provided buffer, random hexamers at a final concentration of 0.01 µg/µL (Thermo Scientific, S0142), dNTPs at a final concentration of 0.5 mM and Ribolock RNase inhibitor (Thermo Scientific, EO0381), following manufacturer’s instructions.

DdPCR was performed using the QX200 Droplet Digital PCR system (Bio-Rad). Each reaction included ddPCR Supermix (no dUTP) (Bio-Rad, Cat # 1863024), forward (AGAGTGGCTCAAACGTTGCT) and reverse (TCCAATGCAGTCGGTAGTGA) primers (IDT), probes (IDT) specific for WT (ATT +G+GA GA+a +GT+C T+CT TG) and Var (C+GA TT+G +GAG A+g+G TCT) templates and diluted cDNA. The symbol “+” denotes the sites where LNA was added to raise the Tm of the probe and the nucleotides represented by a lower case letter in the probe sequences highlight the site of point mutation that distinguishes Var sequence from WT. Final concentration of each primer and probe was 0.5 µM. Droplets were generated according to the manufacturer’s instructions and transferred to a 96-well PCR plate. Plates were heat sealed, and targets were amplified using the following thermocycling conditions, with all ramp rates set to 2°C/s: 95°C, 10 min; [94°C, 30 s; 58°C, 1 min] x 49; 98°C, 10 min; 4°C, hold. Data were analyzed in QuantaSoft software (Bio-rad, Version 1.7.4.0917), and fluorescence thresholds were manually set based on negative controls.

### Plaque assay to determine infectious titers and to isolate plaques

Overlay for plaque assays comprised a 1:1 mixture of complete plaque assay medium (2X Medium 199, 5% FBS, 1% L-glutamine, 1% Penicillin-streptomycin, 1% amphotericin B) and 2% Difco-Bacto agar (BD Diagnostics).

L929 cells were seeded at a concentration of 2×10^6^ cells/well in 6-well plates. Cell culture media was removed and 200 µL of aggregated or non-aggregated forms of T3D reovirus was added (no prior wash step). After a 1 h attachment period at room temperature, inoculum was removed and cells were washed once with DPBS (with Calcium and Magnesium; Corning) (81). Three milliliters of overlay were added, and plates were incubated at 37°C for 6 days. Cells were supplemented with 2 mL of overlay at 3 days post-infection. On day 6, cells were stained through addition of 1% neutral red (Thermo Scientific) solution prepared in a 1:1 mixture of complete media and agar and incubated at 37°C for 18 h.

Plaque isolates were harvested by identifying well-separated viral plaques (at least 1 cm apart) and excising a plug of overlay over each using a sterile 1 mL serological pipette. Each plug was dispensed into 160 µL PBS.

### Genotyping viral plaques using probe-based ddPCR

RNA was extracted from plaque isolates using either the Quick RNA 96 extraction kit (Zymo # R1035) or the NucleoMag RNA extraction kit (Macherey-Nagel # 744350.4), following manufacturer’s instructions. NucleoMag RNA extraction was performed with assistance from Eppendorf’s epMotion 5075 liquid handler. RNA was eluted in either 40 µL (Zymo) or 60 µL (NucleoMag) of nuclease-free water. 12.8 µL of RNA was used for cDNA synthesis with Maxima RT, as described above.

To determine whether individual plaques were derived from WT, Var or a mixture of both viruses’ cDNA was diluted 1:20 in nuclease free water and then 6 µL was used to prepare ddPCR reactions as outlined above.

Genotype assignment was based on probe-specific detection of WT or Var alleles according to probe-specific fluorescence (FAM for WT and HEX for Var), enabling discrimination of WT, Var, double-positive and negative droplets. Probe specificity was validated, confirming that each probe bound to its target without minimal off-target binding. The limit of detection for this assay was set to 787.5 genome copies (GC)/µL based on background signal observed for WT probe on Var template and Var probe on WT template.

### Measurement of co-infection and reassortment

L929 cells (5.5×10^5^ cells/well in 12-well plates) were inoculated with 200 µL of aggregated or non-aggregated WT-Var mixtures at MOIs of 1.0 or 0.10 genome copies per cell. After a 1 h attachment period at 37°C, inoculum was removed, cells were washed once with DPBS (with calcium and magnesium), 1 mL of complete JMEM was added to each well, and plates were incubated at 37°C for 24 h.

Flow cytometry to analyze the proportion of cells infected: At the end of the 24 h incubation, cells were detached from the plate using trypsin (Corning) and moved to a 96-well v-bottom plate for immunostaining. Cells were washed with FACS buffer (2% FBS in 1X PBS) and then resuspended with Zombie NIR dye (BioLegend) to stain dead cells. Next, cells were incubated for 15 min with 0.1 µg/mL rat anti-mouse CD16/CD32 Fc block (BD Pharminogen, Clone 2.4G2) at 4°C, followed by fixation and permeabilization according to the BD Cytofix/Cytoperm kit (Bd biosciences # 554714). Cells were then stained with 0.31 mg/mL mouse monoclonal anti-σ3 antibody (clone 10C1) for 45 minutes at 4°C and then washed twice with 1X BD Perm/Wash buffer before adding an AlexaFluor-647 conjugated donkey anti-mouse secondary antibody (Invitrogen A31571) at a 1:1000 dilution. All samples were analyzed on BD LSRFortessa X-20 instrument at a fixed flowrate of 12 µL/min. Compensation was performed using UltraComp beads (Invitrogen, Cat # 01-2222-42). Data were analyzed using FlowJo 10.9 software.

Genotyping viral plaques by high-resolution melt (HRM) analysis: To distinguish WT and Var segments and thereby quantify reassortment, we applied HRM analysis to plaque isolates. At the end of the 24 h incubation, co-infected cell cultures were subjected to three freeze-thaw cycles at -80°C to disrupt cells and release viruses. Plaque assays were then performed using the cell lysates and plaques were isolated. RNA was extracted from plaque isolates and converted to cDNA using Maxima RT as described above. cDNA was diluted 1:3.5 by adding 50 µL of nuclease-free water to 20 µL of cDNA. The diluted cDNA was used as input for qPCR with 0.4 µM mixture of segment-specific primers as previously described (63). 5 µL reactions were prepared in 384-well plates using Precision Melt Supermix (Bio-Rad) and run on a CFX384 Touch Real-Time PCR Detection System (Bio-Rad). Amplification was assessed using CFX Manager 3.1 software and genotypes were assigned using Precision Melt Analysis 1.3 software (Bio-Rad) based on the distinct melting curves of WT and Var templates. Plaques carrying any combination of WT and Var gene segments were defined as reassortant and % reassortment was calculated using Equation 2.

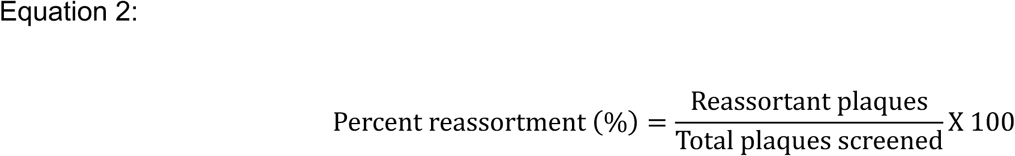

### Data analysis and figure preparation

All statistical analyses were performed using GraphPad Prism 10 version 10.4.2. Figures were created using GraphPad Prism 10, BioRender and Adobe Illustrator.

## Acknowledgments

We thank Dr. Ted Whitworth from Emory University for excellent technical assistance with the preparation of resin-embedded virus infected cell samples and their imaging by electron microscopy. This project was funded by the National Institutes of Health (NIH) through R01 AI146260. Additional support and training were provided by the NIH through T32 AI138952 and the Infectious Disease Across Scales Training Program (IDASTP) of Emory University. We would also like to thank Dr. Elizabeth Draganova and her lab at Emory University for access to their Zetastar dynamic light scattering (DLS) instrument (supported by NIH DP2 award GM154151), which was used for data acquisition and analysis in this study. Furthermore, this research was supported in part by Emory University’s Robert P. Apkarian Integrated Electron Microscopy Core Facility (RRID: SCR_023537), which is subsidized by the School of Medicine and the Emory College of Arts and Sciences, and by the Emory Flow Cytometry Core (EFCC), one of the Emory Integrated Core Facilities (EICF), subsidized by the Emory University School of Medicine. Additional support was provided by the Georgia Clinical & Translational Science Alliance of the National Institutes of Health under award number UL1TR000454 and UL1TR002378. The content is solely the responsibility of the authors and does not necessarily reflect the official views of the National Institutes of Health.

---

**Supplementary Figure 1:**
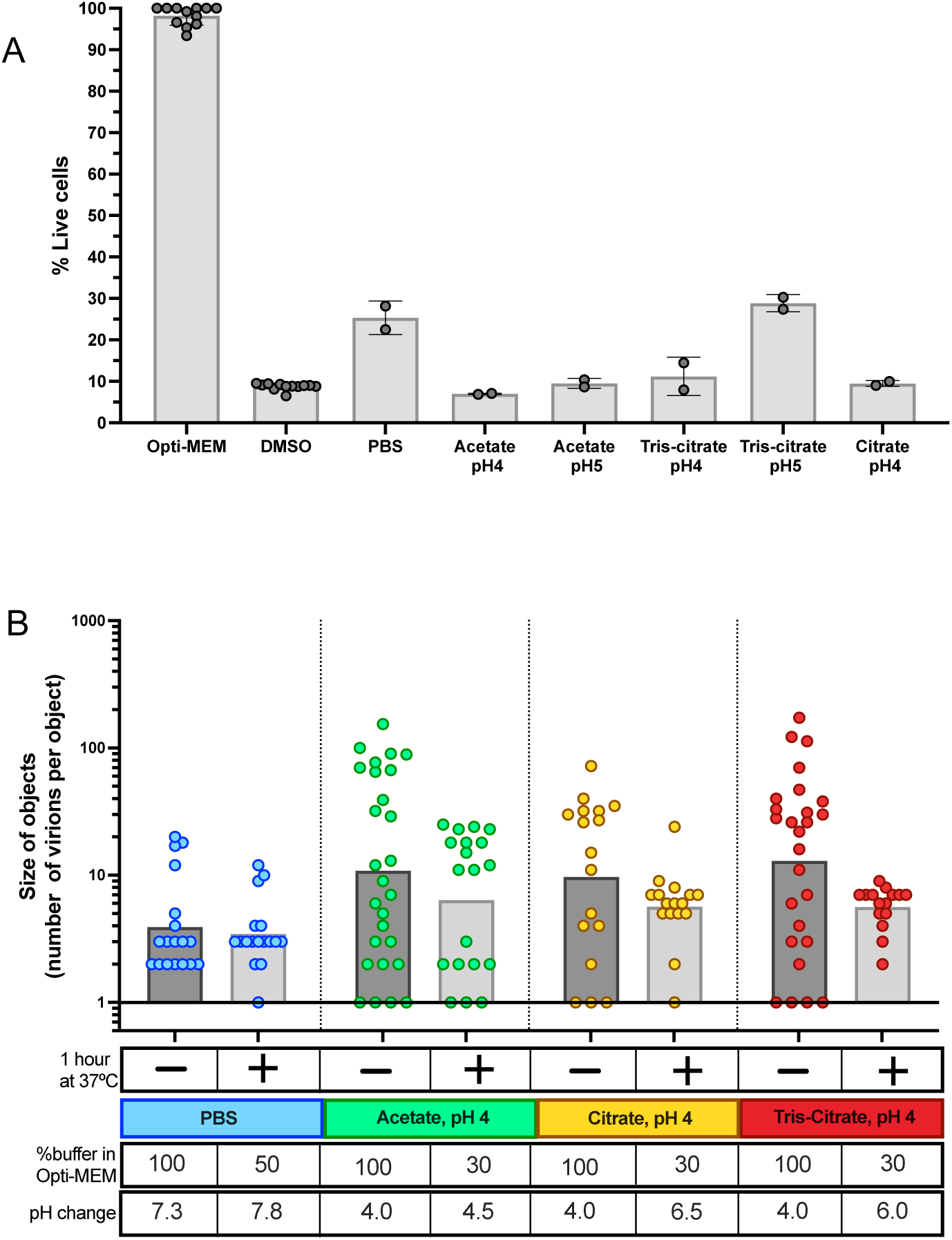
Low pH buffers and PBS are cytotoxic to L929 cells, and reovirus aggregation is reversible and sensitive to buffer pH. **(A)** Viability of L929 cells was measured in culture media (Opti-MEM), DMSO, PBS and in various low pH buffers. Live cell percentage was calculated by normalizing the absorbance reading for each condition to that of the Opti-MEM control. DMSO was included as a positive control for cell death. Data represents 2 biological replicates, each with 3 technical replicates. Bars represent the mean ± s.e.m. A Kruskal-Wallis test indicated significant differences across groups (***p=0.0001). Pairwise comparisons to Opti-MEM were assessed using two-sided Mann-Whitney U tests, *p<0.05. **(B)** Quantification of aggregation levels was measured by dynamic light scattering (DLS), based on the size of viral objects. Treatment conditions are indicated in the table under the plot. Data were pooled from 3 biological replicates. Each data point represents a viral object of size ≥ 1. Bars represent the geometric mean. ** p < 0.003; * p < 0.03; ns, non-significant by unpaired Kolmogorov-Smirnov nonparametric test.

**Supplementary Figure 2:**
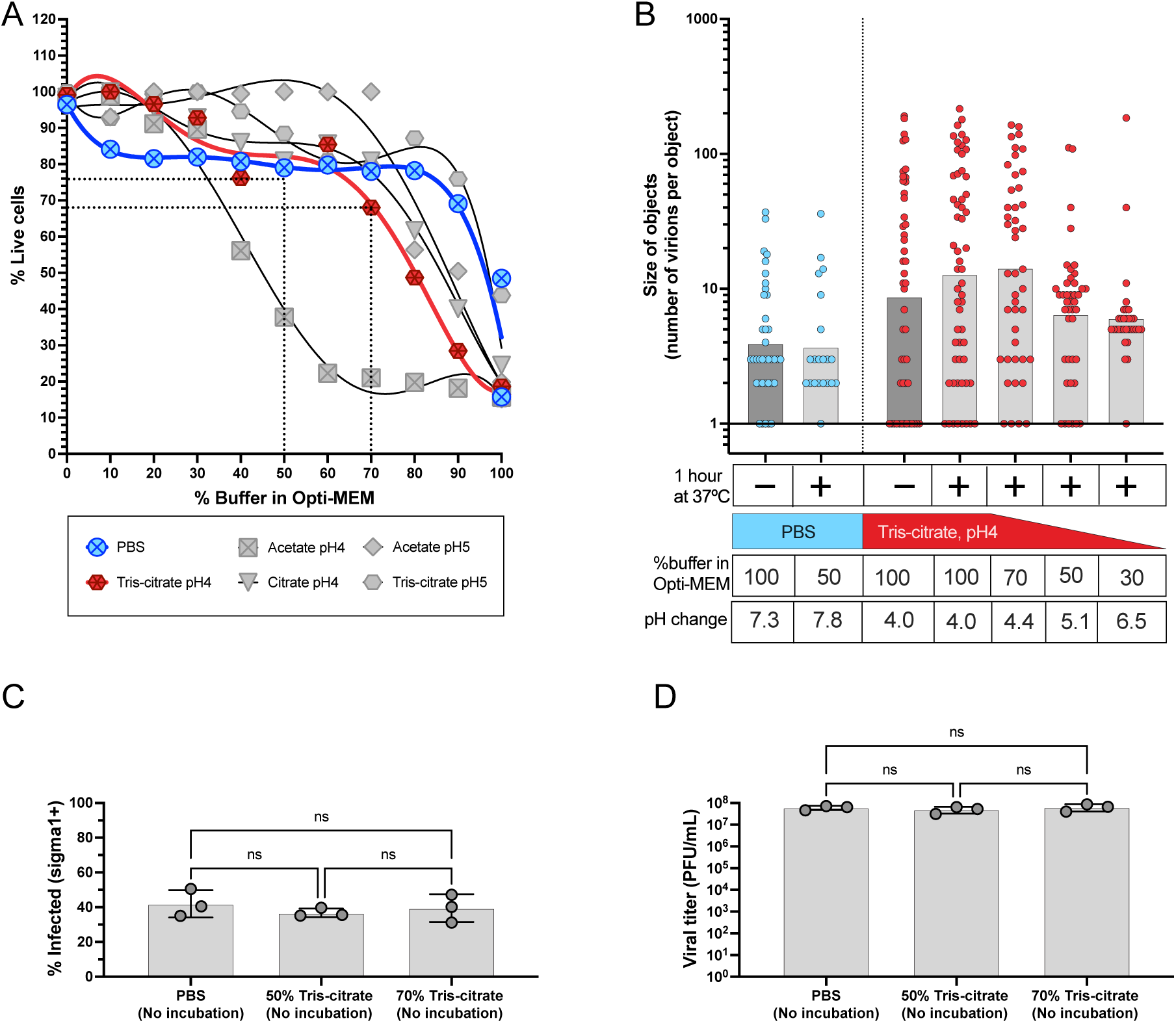
Optimization of aggregation conditions to reduce cellular cytotoxicity. **(A)** Viability of L929 cells following exposure to PBS and low pH buffers diluted with varying proportions of Opti-MEM culture media, ranging from 10% to 100%. Data represent the mean of 2 biological replicates, each with 2 technical replicates. Nonlinear curve fitting was performed using a fifth-order polynomial regression model. Curves were evaluated using goodness-of-fit (R^2^) values (for all, R^2^>0.7, Sys.x <13.64). **(B)** Quantification of aggregation levels measured by DLS, based on the size of viral objects. Data represents 2 biological replicates, each with 3 technical replicates. Each data point represents a viral object of size ≥ 1. Bars represent the geometric mean. **p < 0.003, *p < 0.03 by unpaired Kolmogorov-Smirnov nonparametric test. **(C-D)** Assessing buffer toxicity and viral infectivity of Opti-MEM-buffer mixtures in the absence of aggregation. Infection was measured by percentage of infected cells using flow cytometry **(C)** and plaque titers from cell lysates **(D)**. Each data point represents a biological replicate. Bars represent the mean ± s.d. ns, non-significant by ordinary one-way ANOVA followed by Tukey’s multiple comparison test.

**Supplementary Figure 3:**
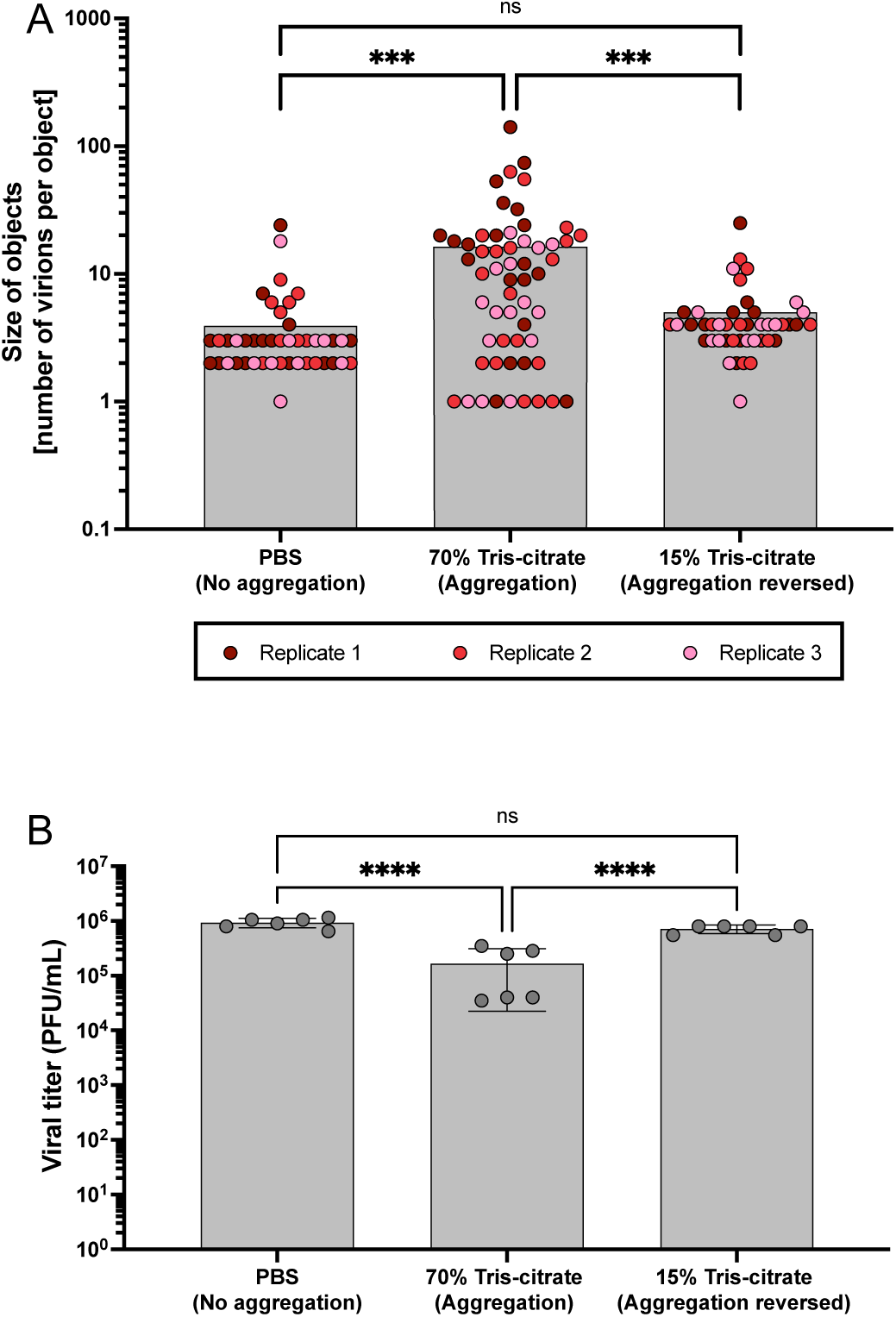
Reovirus is stable in low pH buffers; buffer induced viral aggregation is reversible and pH dependent. **(A)** Quantification of aggregation levels was performed by measuring the size of viral objects using DLS following treatment with Opti-MEM. Each data point represents an individual viral object of size ≥ 1. Data is pooled from 3 biological replicates. Bars indicate mean. ****p < 0.0001; ***p < 0.001; ns, not significant by mixed-effects model followed by Tukey’s multiple comparison test. **(B)** Infectious titers of virus preparations following treatment with Opti-MEM. Each data point represents an independent biological replicate. Bars represent the mean ± s.d. ****p < 0.0001; ns, not significant by ordinary one-way ANOVA followed by Tukey’s multiple comparison test.

**Supplementary Figure 4:**
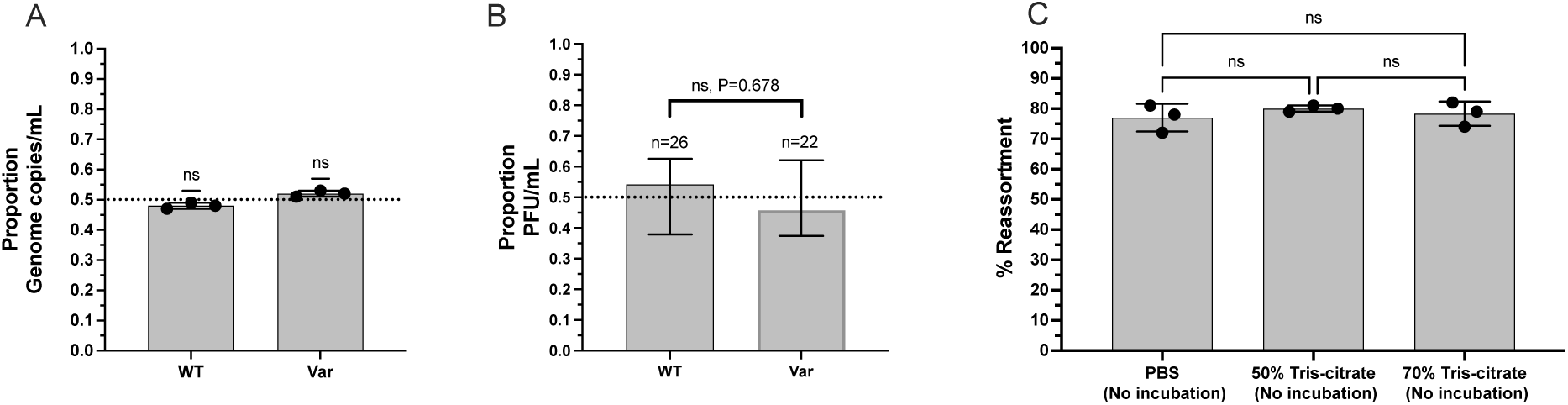
Validation of 1:1 WT-Var mixture and PBS as a negative control for co-infection assays. **(A)** Validation of 1:1 WT and Var viral mixtures: Genome copy ratios were quantified across 3 biological replicates. Bars represent mean ± 95% confidence interval. Each data point represents a biological replicate. Statistical comparison to the expected proportion of 0.5 was performed by one-sample t-test. ns, non-significant. **(B)** Infectious viral particle ratios were validated by genotyping 48 plaques derived from the mixture of WT and Var viruses. Bars represent mean ± 95% confidence interval. A two-sided exact binomial test was used to determine whether the proportions of WT and Var differ significantly (p=0.678) and 95% confidence intervals were calculated using Clopper-Pearson method. **(C)** Reassortment frequency in L929 cells co-infected with WT and Var reoviruses prepared in the indicated buffers without incubation to induce aggregation. MOI was 1 GC/cell. Data shown are mean ± s.d. and represent 3 biological replicates. Each data point represents the percentage of reassortant plaques out of 32 plaques. ns, not significant by ordinary one-way ANOVA followed by Tukey’s multiple comparison test.

